## Supplementary Information for "A Versatile Oblique Plane Microscope for Large-Scale and High-Resolution Imaging of Subcellular Dynamics"

| **Figure** | **Sample** | **Label** | **Voxel Size (nm^3^)** | **Volumetric Imaging Rate (Hz)** | **Post-Processing** | **Rendering Software** |
| --- | --- | --- | --- | --- | --- | --- |
| 2D | 100 nm beads | Fluoresbrite YG | 57.5x57.5x100 | N/A | Deconvolution | Fiji |
| 2H | 100 nm beads | Fluoresbrite YG | 14.375x14.375x14.375 | N/A | Raw, or deconvolved | Fiji |
| 2I | MV3 | GEMS-T-Sapphire | 57.5x57.5x115 | N/A | Deconvolution | Fiji |
| 3A | U2OS | EGFP-Sec61b | 115x115x 250 | N/A | Deconvolution + Histogram Matching | Imaris + Fiji |
| 3B-C | RPE hTERT | mEmerald-Vimentin | 115x115x100 | N/A | Deconvolution | Fiji |
| 3D-E | ARPE | EGFP-AP2 | 115x115x100 | N/A | Deconvolution + Histogram Matching | Fiji |
| 3F-H | MV3 | CAAX-HALO | 115x115x250 | N/A | Deconvolution | Fiji |
| 4 | NK-92 + K562 | LifeAct-mScarlet & LCK-mVenus | 115x115x 250 | N/A | Deconvolution + Histogram Matching | Fiji |
| 5 | 1205Lu | meGFP-α-Tubulin & 3xNLS-mScarlet-i | 115x115x400 | N/A | Deconvolution | Fiji |
| 6A | Cardiomyocytes | Fluo-3 AM | 115x115x3000 | N/A | Deconvolution+ Histogram Matching | Imaris |
| 6B | MV3 | GEMs-T-Sapphire | 115x115x500 | N/A | Deconvolution+ Histogram Matching | MATLAB |
| 7A-D | MEFs | PA-Rac1 & mCherry | 115x115x230 | N/A | Simple Ratio bleach correction | Fiji+ Imaris + MATLAB |
| 8A | Cortical Neurons | GCaMP6f | 115x115x460 | N/A | Deconvolution |  |
| 8B | Drosophila m. | Gap43-mCherry | 115x115x200 | N/A | Deconvolution |  |
| 9A-C | Mouse brain slice | Nuclei-DAPI | 115x115x100 | N/A | Flat-field, stitching, fusion | Fiji |
| 9D-G | Human lung | Nuclei-DAPI, SFTPC-Alexa594, & ACE2-mRNA-Alexa647 | 115x115x100 | N/A | Flat-field, stitching, fusion | Fiji + 3D-script |
| 1-S3 | U2OS | EGFP-Tractin | 115x115x100 | N/A | Raw data | Fiji |
| 2-S1 | 100 nm beads | Fluoresbrite YG | 0.068x0.068.x0.068 nm^-1^ | N/A | Rotationally averaged. | MATLAB |
| 2-S2 | 100 nm beads | Fluoresbrite YG | 115x115x115 | N/A | Raw data | Fiji |
| 2-S3-LLSM | 100 nm beads | FluoSpheres | 115x115x115 | N/A | Raw data | Fiji |
| 2-S3-SD | 100 nm beads | Fluoresbrite YG | 115x115x115 | N/A | Raw data | Fiji |
| 2-S3-OPM | 100 nm beads | Fluoresbrite YG | 115x115x115 | N/A | Raw data | Fiji |
| 2-S4 | 100 nm beads | Fluoresbrite YG | 115x115x115 | N/A | Raw data | Fiji |
| 2-S5 | 100 nm beads | Fluoresbrite YG | 115x115x115 | N/A | Raw data | MATLAB |
| 3-S1 | U2OS | EGFP-Sec61b | 115x115x 250 | 1.19 | Raw data | Fiji |
| 3-S2 | MV3 | CAAX-HALO | 115x115x250 | 0.917 | Deconvolution |  |
| 8-S1 | Drosophila m. | Gap43-mCherry | 115x115x200 | .043 | Deconvolution | Fiji |
| M1 | U2OS | EGFP-Sec61b | 115x115x 250 | 1.19 | Deconvolution + Histogram Matching | Imaris |
| M2 | RPE hTERT | mEmerald-Vimentin | 115x115x100 |  | Deconvolution | Fiji |
| M3 | ARPE | EGFP-AP2 | 115x115x100 | 0.4 | Deconvolution + Histogram Matching | Fiji |
| M4 | ARPE | EGFP-AP2 | 115x115x100 | 0.4 | Deconvolution + Histogram Matching | Fiji |
| M5 | MV3 | CAAX-HALO | 115x115x250 | 0.91 | Deconvolution | Fiji |
| M6 | NK-92 + K562 | LifeAct-mScarlet & LCK-mVenus | 115x115x 250 | 0.09 | Deconvolution + Histogram Matching | Fiji |
| M7 | 1205Lu | EGFP-Tubulin & NLS-mScarlet | 115x115x400 | .05 | Deconvolution | Fiji |
| M8 | Cardiomyocytes | Fluo-3 AM | 115x115x500 | 10.4 | Deconvolution+ Histogram Matching | Imaris |
| M9 | MV3 | GEMs-T-Sapphire | 115x115x500 | 13.7 | Deconvolution+ Histogram Matching | MATLAB |
| M10 | N/A | N/A | N/A | N/A | N/A | Mathematica |
| M11 | MEFs | PA-Rac1-mCherry & mCherry | 115x115x230 | 0.1 | Simple Ratio bleach correction | Fiji+ Imaris |
| M12 | MEFs | PA-Rac1-mCherry & mCherry | 115x115x230 | 0.1 | Simple Ratio bleach correction | Fiji+ Imaris |
| M13 | Cortical Neurons | GCaMP6f | 115x115x920 | 7 | N/A | Fiji |
| M14 | Drosophila m. | Gap43-mCherry | 115x115x200 | .043 | N/A | Fiji |
| M15 | Mouse brain slice | Nuclei-DAPI | 115x115x100 | N/A | Flat-field, stitching, fusion. | Fiji |
| M16 | Human lung | Nuclei-DAPI, SFTPC-Alexa594, & ACE2-mRNA-Alexa647 | 115x115x100 | N/A | Flat-field, stitching, fusion, deconvolution | Fiji + 3D-script |

**Supplementary File 1**. Description of sample type, fluorescent labels, imaging conditions, data post-processing, and rendering, for each figure.

| **Probe names** | **Gene specific sequence - detection probe specific sequence** |
| --- | --- |
| hsHR2X-ACE2-1719 | aatgctagggtccagggttc TTATACGTCGAGTTGAACGTCGTAACA |
| hsHL2X-ACE2-1719 | TAGCGCTAACAACTTACGTCGTTATG tgattttccaagcctcagca |
| hsHR2X-ACE2-58 | agcagttacagcaacaaggc TTATACGTCGAGTTGAACGTCGTAACA |
| hsHL2X-ACE2-58 | TAGCGCTAACAACTTACGTCGTTATG tgagaaggagccaggaagag |
| hsHR2X-ACE2-1457 | ctcgcttcatctcccaccac TTATACGTCGAGTTGAACGTCGTAACA |
| hsHL2X-ACE2-1457 | TAGCGCTAACAACTTACGTCGTTATG tttttcatccactggtcttt |
| hsHR2X-ACE2-2872 | atgcatgccattctcaatcc TTATACGTCGAGTTGAACGTCGTAACA |
| hsHL2X-ACE2-2872 | TAGCGCTAACAACTTACGTCGTTATGttgcagctacaccagttccc |
| hsHR2X-ACE2-1038 | actgctttctgaacatttcc TTATACGTCGAGTTGAACGTCGTAACA |
| hsHL2X-ACE2-1038 | TAGCGCTAACAACTTACGTCGTTATGtgggtccgttagcatggaat |

**Supplementary File 2.** Encoding probe sequences for proximity ligation RNA fluorescence *in situ* hybridization.

| **Cell Line** | **Source** | **Authentication** | **Testing** |
| --- | --- | --- | --- |
| MV3 | Friedl Lab | N/A | Mycoplasma negative |
| U2OS | ATCC (HTB-96) | N/A | Mycoplasma negative |
| RPE hTERT | ATCC (CRL-4000) | N/A | Mycoplasma negative |
| ARPE | ATCC (CRL-2302) | N/A | Mycoplasma negative |
| NK-92 | ATCC (CRL-2407) | N/A | Mycoplasma negative |
| K562 | ATCC (CCL-243) | N/A | Mycoplasma negative |
| 1205Lu | Wistar | STR Fingerprint | Mycoplasma negative |
| MEFs | ATCC (SCRC-1040) | N/A | Mycoplasma negative |

**Supplementary File 3.** Source, authentication method, and routine testing performed on cell lines.
